# Divulging diazotrophic bacterial community structure in Kuwait desert ecosystems and their N2-fixation potential

**DOI:** 10.1101/712265

**Authors:** M. K. Suleiman, A. M. Quoreshi, N. R. Bhat, A. J. Manuvel, M. T. Sivadasan

## Abstract

Kuwait is a semi-arid region with harsh climatic conditions with poor available soil nutrient essential for the growth of plants. Kuwait’s ecosystem is relatively N-poor ecosystem when compared to the other ecosystems. Biological nitrogen fixation (BNF) is a spontaneous process in which diazotrophic bacteria fixes the atmospheric nitrogen directly in to the bionetwork. At present, there is limited information on free-living and root associated nitrogen-fixing bacteria, their potential to fix nitrogen to aid natural plant communities in the desert of Kuwait. In this study, free-living N_2_-fixing bacteria were enriched and isolated from the rhizospheric soil of three keystone native plant species of Kuwait; such as *Rhanterium epapposum, Farsetia aegyptia*, and *Haloxylon salicornicum*. Root associated bacteria were directly isolated from the root nodules of *Vachellia pachyceras*. In this study, a number of free-living and root associated dizotrophs were isolated from various rhizospheric soils of three native shrubs and root nodules from one tree species. The screened isolates were assessed for nitrogen-fixing ability and identified using Acetylene Reduction Assay (ARA) and 16s rRNA gene sequencing, respectively. Our study successfully identified all the 50 nitrogen-fixers isolated initially and out of that, 78% were confirmed as nitrogen-fixers using ARA. Among the identified nitrogen fixers, the genus *Rhizobium* is dominant in rhizospheric soil of *Rhanterium epapposum*, whereas *Pseudomonas* and *Rhizobium* are dominant in the rhizospheric soil of *Farsetia aegyptia*, and *Haloxylon salicornicum* respectively. The species *Agrobacterium tumefaciens* is found dominant in the root nodules of *V. pachyceras*. The current results indicate that plant species and their rhizospheric effects are important drivers for specificity of microbial diversity in arid soils. To our knowledge, this study is the first investigation of culture-based isolation, molecular identification, and evaluation of N_2_-fixing ability of diazotrophs from Kuwait desert environment.

## Introduction

Symbiotic bacterial association with plants mediates the most of the nitrogen fixation process in terrestrial ecosystems. Nevertheless, non-symbiotic N_2_ fixation by free-living nitrogen-fixing bacteria harboring in soil systems can considerably contribute to the nitrogen fixation pool in various ecosystems, particularly in desert ecosystems [1, 2]. Nitrogen is an essential element required for plant growth and development, considered a limiting factor in plant productivity [3, 4] and can affect life of microbes and other living organisms [5]. Nitrogen is an important component of amino acids, the building blocks of proteins; chlorophyll, the green pigment required for photosynthesis; ATP, the primary energy carrier; nucleic acid, the genetic material of living organisms [6]. Thus, nitrogen plays a fundamental role in the growth and productivity of crops. Nitrogen is abundant in the atmosphere (almost 80%) as molecular N_2_ but cannot be utilized directly by the plants. Therefore, biological N_2_ fixation is necessary to convert the elemental N_2_ into ammonia, which is readily available to bacteria and plants. Nitrogen fixation is an essential step in the global nitrogen cycle as it restores and recompenses the overall nitrogen lost because of denitrification [7]. The atmospheric nitrogen is fixed into the soil, utilized by the plants, and returned to the soil and atmosphere through a process called nitrogen cycle. In general, nitrogen cycle is a four-step process and comprises of, nitrogen fixation (conversion of atmospheric dinitrogen to nitrate or ammonia), nitrification (conversion of ammonia to nitrate ions), assimilation (incorporation of the nitrates in to the plant tissues), and denitrification (conversion of nitrate present in the dead plants to dinitrogen).

Diazotrophs are a specialized group of bacteria possessing nitrogenase enzyme system and capable of biological nitrogen fixation, in which atmospheric di-nitrogen (N_2_) converts into readily available form of fixed nitrogen in the biosphere [1]. The diazotrophs can live freely in the soil (example: *Pseudomonas, Azotobacter, Cyanobacteria*) or can establish symbiotic association with certain group of plant species (example: *Rhizobium, Bradyrhizobium*). These diazotrophs or Biological Nitrogen Fixers associated with the plants and rhizosphere are possible alternative for inorganic nitrogen fertilizer that supports the growth and productivity of the plants and sustainability of soil [8]. Inoculation of BNF increased nitrogen fixation in *Phaseolus vulgaris* L. [9]. In addition to this, it was reported similar growth of *Oryza sative* L (rice crop) when inoculated with plant growth promoting rhizobacteria (PGPR) plus half fertilization and full fertilization without inoculation [10]. It was also suggested that the use of P-solubilizing diazotrophs is a good strategy to promote P solubilization and / or N use efficiency in rice plants [11]. Although diazotrophs are one of the key performer for plant available nitrogen source, it is poorly understood about their communities associated with soil, rhizosphere, endosphere, and their importance in arid ecosystems [1, 12].

Arid regions are one of the harshest regions on this planet and comprise huge challenges to maintain vegetation growth and plant productivity. The available nutrients for native plant are deficient in Kuwait’s desert soil due to lack in organic matter (<1%), low clay materials, high in calcareous materials, and low in essential nutrients [13]. Availability of soil nutrients is one of the important factors for successful restoration of the degraded soil. Microorganisms and their bioactivity could play an essential role in nutrient cycling and improvement of soil fertility in arid terrestrial ecosystems. Despite prevailing severe environmental conditions in desert, a broad diversity of organisms, comprising plants have adapted to extreme conditions, such as low rainfall, extremely high temperatures, high level of solar radiation, very low nutrients level, and high salinity by developing different adapting mechanisms [14]. It is believed that desert microbiome may be a key factor by which desert plants adapt to these extreme conditions [15–17]. Microbes present in desert ecosystems are believed to be proficient in enhancing plant growth and stress tolerance, and play essential roles in nutrient cycling [18, 19]. Therefore, enhancement of arid lands soil nutritional conditions requires a prudent management of soil resources for sustainable productivity. The use of beneficial bio-inoculants could be one of the approaches for desert ecosystems. Therefore, there is a necessity to identify N_2_ fixing bacteria that may provide as bio inoculant for an alternative to inorganic chemical fertilizers for sustainable desert plant growth and development. The approach is considered for an environmentally safe strategy to improve the quality of the soil without introducing chemical fertilizers. To our knowledge, there is no data existed on indigenous nitrogen-fixing bacterial communities from Kuwait’s desert soils and the details on their nitrogen-fixing ability. This study reports for the first time on the isolation and characterization of free-living as well as rhizobacterial communities associated with rhizospheric soils of selected native shrubs and roots of *Vachellia pachycras* from Kuwait’s desert.

The aim of this study was to isolate, screen, and identify the free-living and root associated diazotrophic communities present in the rhizospheric soil of Kuwait’s economically important native plants, *Rhanterium epapposum, Farsetia aegyptia*, and *Haloxylon salicornicum* and root nodules of *Vachellia pachycras*. Furthermore, an effort was undertaken to evaluate nitrogen-fixing ability of the isolated bacterial strains by the Acetylene Reduction Assay (ARA).

## Materials and methods

### Sampling

The rhizospheric zone (soil around the plant root zone) of the native plants of Kuwait such as *Rhanterium epapposum, Farsetia aegyptia*, and *Haloxylon salicornicum* located at KISR’s Station for Research and Innovation (KSRI; GPS: N 29° 09.904’; E 047° 41.211’), Sulaibiya were selected for the isolation of free-living nitrogen fixing bacteria. Triplicate rhizosphere soil samples were collected from the mentioned plant species by uprooting the root system and were transferred to sterile zip lock bags. Similarly, triplicate root samples with root nodule were collected from *Vachellia pachyceras* found at different locations inside KSRI, Sulaibiya and transferred to sterile zip lock bags. The collected soil and root samples were transported to the laboratory on ice in a cooler box.

### Isolation and primary screening of diazotrophic bacteria

An enrichment culture method using of nitrogen-free semi-solid malate media (NDSM) and nitrogen deficient malate media (NDMM) with bromothimol blue indicator [20] was adopted for the isolation of putative free-living nitrogen-fixing bacteria from the 1g of rhizospheric soil of *Rhanterium epapposum, Farsetia aegyptia*, and *Haloxylon salicornicum*. Whereas, a modified procedure was followed for the isolation of *Rhizobium* on yeast manitol agar (YMA) medium containing 0.0025% Congo red from 100mg of the root nodules of *Vachellia pachyceras* [21]. Root nodules were sterilized using 95% (v/v) ethanol for 10 s and washed at least 7 times with sterilized distilled water. The plates were incubated at 28°C for 3 to 7 days, and single colonies of putative N_2_-fixers were carefully selected and restreaked on YMA agar media for further purification. The procedure was repeated and the pure culture of all putative N_2_-fixers were stocked in YMB with 15% glycerol and stored at -80° C freezer. Slants were prepared for the pure cultures and kept in the refrigerator for day-to-day experiments. All the bacterial isolates were labelled with the first three letters of the plant species-Source of isolation-Serial number (e.g. Rha-S-1 [*Rhanterium epapposum*-Soil-Bacteria No.1]).

Primary screening for potential N_2_-fixers of the pure bacterial cultures was determined by measuring the blue color zone formed by the isolates when grown on nitrogen-deficient malate Media (NDMM) with bromothymol blue. The color changes to blue color was considered as an indicator for positive effect of nitrogen-fixers [22].

### Biochemical identification of the isolates

Biolog® Gen III Microbial Identification System (Biolog Inc., Hayward, CA, USA) was used for the biochemical identification of selected putative N_2_-fixers isolates from the rhizospheric soil and root nodule of test species. The system identifies the microorganisms based on their ability to metabolize all major classes of biochemical compounds, in addition to determining other important physiological properties such as pH, salt and lactic acid tolerance, reducing power, and chemical sensitivity. Pure bacterial isolates were streaked on BUG medium and incubated for 24 hr at 30°C. At the end of the incubation period, individual colonies were suspended in IFA inoculating fluid to a cell density 90-98% as recommended by the Biolog Inc. One hundred microliter of the prepared suspension were transferred to each well of the 96-well microplate, and the plates were incubated in Omnilog incubator at 33°C for 24 hr. Colorimetric responses are measured using the Biolog-supplied Omnilog and compared with Biolog® database for identity conformation (Gen III database and characteristics v2.7).

### Acetylene reduction assay (ARA)

Bacterial strains were grown in modified Fraquil medium, a N free medium developed based on Fraquil medium and contains 5 × 10^−3^ M KH_2_ PO _4_, 2.3 × 10^−3^ M K_2_ HPO_4_, 6.8 × 10^−4^ M CaCl_2_, 4.05 × 10^−4^ M MgSO_4_ (7H_2_O), 10^−8^ M CuCl_2_ (2H_2_ O), 2.25 × 10^−7^ M MnCl_2_ 4(H_2_O), 2.43 × 10^−8^ M CoCl_2_ (6H_2_O), 5.3 × 10^−8^ M ZnSO_4_ (7H_2_O), 5 × 10^−7^ M MoO_4_Na_2_, 5 × 10^−6^ M FeCl _3_ (7H_2_O), 10^−4^ M EDTA, 5.5 × 10^−2^ M Glucose, 5.48 × 10^−2^ M D-Mannitol, pH 6.7. Bacteria were inoculated into the modified Fraquil medium with 3% agar using sterile and disposable inoculation spreaders and incubated at 28°C in a thermoregulated incubator (infor, multitron).

The nitrogen-fixing ability of all the isolates found positive in the primary screening was determined by ARA. In this procedure, ethylen produced by the bacterial culture was quantified and the results were expressed in nano moles of C_2_H_4_ produced h^-1^ culture^-^1. Bacteria grown in petri dishes were collected after 1, 3 and 7 d of incubation. Dishes were transferred to 250 mL glass jars. The jars were closed with a lid equipped with a septum for gas sampling. Fifteen percent of the headspace was replaced by acetylene (C_2_H_2_). Acetylene was produced by the reaction of calcium carbide with water (equation: CaC_2_ + 2H_2_O Ca (OH)_2_ + C_2_ H_2_) in a tedlar bag. Petri dishes were incubated in the presence of C_2_H_2_ for 6 h. Three milliliters of Headspace were collected after three and six hours of incubation, transferred into vacuumed three-milliliter vials. Ethylene concentration was then qualified by gas chromatography using Shimatzu 8A (Shimatzu, Japan) equipped with a flame ionisation detector (GC-FID). A control N_2_ fixer *Azotobacter vinelandii*, was grown and assayed under similar conditions for comparison. Apart from the modified Fraquil agar, the test bacterial strains were grown in Yeast Mannitol broth and agar as well as in modified Fraquil broth and assayed for acetylene reduction under similar conditions.

After completion of the acetylene reduction assay (on day 1, 3 and 7), the bacterial density was estimated for each petridish to normalize ARA data to cell density (OD). This allows a better comparison of N_2_ fixation potential between species. Cells were suspended by adding 2 mL of culture medium to the dishes and disturbing bacterial mats using an inoculation spreader. The solution was then transferred to a 15-mL tube. The operation was repeated until no visible bacterial mats remain at the surface of the dish. Cells were then pelleted by centrifugation and the supernatant removed. Cells were then resuspended in 1 mL of medium and OD (620 nm) measure by UV-Vis spectrometry. Cell suspension were diluted using the culture medium when needed (saturation of the UV detector).

### Molecular identification for the isolates

Aliquots from the fresh culture were scratched with a sterile transfer pipet and mixed in 200 uL of 45 mg/ml lysozyme solution (no. cat. L4919 – Millipore-Sigma Darmstadt, Germany). Suspension of cells was incubated at 37°C for 30 min as recommended by the manufacturer. Concentration of eluted DNA was quantified with a Nanodrop 2000 (Thermo Fisher, Waltham, MA, USA). The polymerase chain amplification reactions (PCR) were performed on the eluted DNA. Amplifications were performed using the bacterial universal primers 358F/907R. PCR conditions were the same using the primers 358F (CTACGGGAGGCAGCAG; [23])/907R (CCGTCAATTCMTTTRAGTTT; [24]). PCR was carried out in a 25-µL reaction and consisted of 1 µL bacterial DNA, 1 U of platinum Taq (Invitrogen), 2.0 mM MgCl_2_ and 0.2 mM dNTP mix. The following thermocycle program was used for amplification: 95°C for 30 seconds followed by 30 cycles of 95°C, 54 °C and 72°C for 50 seconds each, and an extension period of 72°C for 5 min using a MJ Research PTC-225 Peltier Thermal Cycler. Amplifications were visualized on a 1.0% agarose gel electrophoresis (TAE), and amplicons were sequenced using the Sanger sequencing method with two 16-capillary genetic analyzers 3130XL (Applied Biosystems).

DNA sequences were edited with BioEdit version 7.0.5 [25, 26]. The BLASTn algorithm [27] was used to query for highly similar sequences with the Bacterial 16S Ribosomal RNA RefSeq Targeted Loci Project. Sequences were aligned with ClustalX version 1.81 [28]. Phylogenetic analyses were performed in order to identify and/ or place unknown sequence in the taxonomic tree of known species. Phylogenetic analyses were conducted using MEGA 7.0.21 software [29, 30]. Firstly, analyses were realized using the Neighbor-Joining (NJ) method [31] using the Kimura 2-parameter method [32] to visualize approximately the tree topology. Secondly, maximum likelihood (ML) method based on the Kimura 2-parameter model [32] was used to compute a final tree. Initial tree(s) for the heuristic search were obtained automatically by applying Neighbor-Joining and BioNJ algorithms to a matrix of pairwise distances estimated using the Maximum Composite Likelihood (MCL) approach, and then selecting the topology with superior log likelihood value. All analyses were followed by a 1000 bootstrap replicates [33]. Bayesian inference of phylogeny was calculated with MrBayes program, assuming a 4 by 4 model and a non-variable substitution rates among sites – gamma rates. Analyses were based on 2 runs of four Markov chain Monte Carlo analyses where 2, 000, 000 generations were generated, burning fraction at 0.5 rate and sampled every 100 generations for a total of 10, 000 trees generated [34].

## Results

### Isolation and primary screening of diazotrophic communities

Free-living nitrogen-fixing diazotrophs were isolated from various rhizospheric soils of three native shrubs and root nodules from one tree species. A total of 9, 13, and 22 morphologically different pure bacterial strains were isolated from the rhizospheric soil of *Rhanterium epapposum, Farsetia aegyptia*, and *Haloxylon salicornicum*, respectively. In contrast, a total of 61 pure cultures with different morphotypes were isolated from the root nodules of *Vachellia pachyceras*. The results indicates that several species of potential free-living nitrogen fixing bacteria may present in Kuwait desert soils around the selected plant species (Table 1).

**Table 1.** Diameter of Zone of Colorization Developed on Nitrogen Free Malate Media and Nitrogen Fixation Potential of the Bacterial Isolates Isolated from Rhizospheric Soil of Selected Native Plants and Root Nodule of *Vachellia pachyceras*

| Isolate ID Number | Diameter of Zone of Colorization (cm) | Relative Nitrogen Fixation Potential (Acetylene Reduction Assay) | Isolate ID Number | Diameter of Zone of Colorization (cm) | Relative Nitrogen Fixation Potential (Acetylene Reduction Assay) |
| --- | --- | --- | --- | --- | --- |
| Rhizospheric Soil of *Rhanterium epapposum* |  |  |  |  |  |
| RhaS-1 | 0.1 | ++++ | RhaS-6 | 0.6 | ++ |
| RhaS-2 | 0.3 | +++ | RhaS-8 | 0.0 | - |
| RhaS-3 | 1.1 | L | RhaS-9 | 0.8 | ++++ |
| RhaS-4 | 0.8 | ++ | RhaS-10 | 0.0 | - |
| RhaS-5 | 0.0 | - | RhaS-12 | 1.0 | +++ |
| Rhizospheric Soil of *Farsetia aegyptia* |  |  |  |  |  |
| FarS-1 | 1.5 | +++ | FarS-8 | 2.7 | ++++ |
| FarS-2 | 2.1 | D | FarS-9 | 1.3 | ++++ |
| FarS-3 | 1.3 | D | FarS-10 | 0.1 | L |
| FarS-4 | 1.6 | D | FarS-11 | 0.0 | - |
| FarS-5 | 1.5 | L | FarS-12 | 0.0 | - |
| FarS-6 | 0.1 | D | FarS-13 | 0.0 | - |
| FarS-7 | 0.1 | D | - | - | - |
| Rhizospheric Soil of *Haloxylon salicornicum* |  |  |  |  |  |
| HalS-1 | 2.2 | ++ | HalS-14 | 2.2 | +++ |
| HalS-2 | 0.0 | - | HalS-15 | 0.0 | - |
| HalS-3 | 2.3 | + | HalS-16 | 1.6 | L |
| HalS-5 | 0.0 | - | HalS-17 | 3.2 | D |
| HalS-6 | 3.0 | L | HalS-18 | 0.0 | - |
| HalS-8 | 2.2 | ++++ | HalS-19 | 0.3 | ++ |
| HalS-9 | 2.4 | ++ | HalS-20 | 3.0 | ++ |
| HalS-10 | 0.0 | - | HalS-21 | 0.0 | +++ |
| HalS-11 | 1.7 | ++++ | HalS-23 | 1.8 | D |
| HalS-12 | 3.2 | D | HalS-24 | 0.5 | ++++ |
| HalS-13 | 0.3 | D | HalS-25 | 1.9 | + |
| Root Nodules of *Vachellia pachyceras* |  |  |  |  |  |
| Ac-1 | 1.8 | + | LTN-4 | 0.0 | - |
| Ac-2 | 0.0 | - | LTN-5 | 0.0 | - |
| Ac-4 | 0.0 | - | LTN-6 | 0.0 | - |
| Ac-6 | 0.0 | - | LTN-7 | 0.0 | - |
| Ac-7 | 0.0 | - | LTN-9 | 0.0 | - |

| **Isolate**<br>**ID**<br>**Number** | **Diameter of**<br>**Zone of**<br>**Colorization**<br>**(cm)** | **Relative Nitrogen**<br>**Fixation Potential**<br>**(Acetylene**<br>**Reduction Assay)** | **Isolate**<br>**ID**<br>**Number** | **Diameter of**<br>**Zone of**<br>**Colorization**<br>**(cm)** | **Relative Nitrogen**<br>**Fixation Potential**<br>**(Acetylene**<br>**Reduction Assay)** |
| --- | --- | --- | --- | --- | --- |
| Ac-9 | 1.6 | + | LTN-10 | 2.7 | +++ |
| Ac-10 | 1.6 | + | LTN-11 | 0.0 | - |
| Ac-11 | 0.0 | - | LTN-12 | 0.0 | - |
| Ac-12 | 2.0 | + | LTN-13 | 0.0 | - |
| Ac-13 | 0.0 | - | LTN-14 | 3.4 | ++ |
| Ac-15 | 0.0 | - | LTN-15 | 0.0 | - |
| Ac-16 | 0.0 | - | LTN-17 | 0.0 | - |
| Ac-17 | 0.0 | - | LTN-20 | 0.0 | - |
| Ac-18 | 0.0 | - | ACN-1 | 1.9 | ++ |
| Ac-20 | 0.0 | - | ACN-2 | 2.8 | ++++ |
| Ac-21 | 0.0 | - | ACN-3 | 1.9 | ++++ |
| Ac-22 | 2.1 | + | ACN-4 | 3.6 | L |
| Ac-25 | 1.2 | ++ | ACN-5 | 3.6 | L |
| Ac-26 | 0.0 | - | ACN-6 | 0.0 | L |
| Ac-30 | 2.1 | + | ACN-8 | 0.0 | - |
| Ac-31 | 0.0 | - | ACN-9 | 3.0 | ++ |
| Ac-33 | 0.0 | - | ACN-10 | 0.0 | ++ |
| Ac-34 | 0.0 | - | ACN-11 | 0.0 | - |

**Table 1. cont**
| Isolate<br>ID<br>Number | Diameter of<br>Zone of<br>Colorization<br>(cm) | Relative Nitrogen<br>Fixation Potential<br>(Acetylene<br>Reduction Assay) | Isolate<br>ID<br>Number | Diameter of<br>Zone of<br>Colorization<br>(cm) | Relative Nitrogen<br>Fixation Potential<br>(Acetylene<br>Reduction Assay) |
| --- | --- | --- | --- | --- | --- |
| Ac-35 | 2.1 | + | ACN-12 | 0.1 | +++ |
| Ac-36 | 0.0 | - | ACN-13 | 0.0 | - |
| Ac-38 | 0.0 | - | ACN-14 | 0.2 | D |
| Ac-39 | 0.1 | ++++ | ACN-14<br>(B) | 0.1 | D |
| Ac-40 | 2.1 | + | ACN-15 | 0.0 | - |
| LTN-1 | 0.8 | ++ | ACN-16 | 0.0 | - |
| LTN-2 | 0.0 | - | ACN-17 | 0.1 | D |
| LTN-3 | 0.0 | - |  |  |  |

-: no activity; +: low maximum activity (below 20 nmol/h/OD); ++: moderate maximum activity (20-50 nmol/h/OD); +++: high activity (50-80 nmol/h/OD); ++++: very high maximum activity (above 80 nmol/h/OD); L: Bacteria lost due to contamination; D: Did not grow in N<sub>2</sub> free media to conduct Acetylene Reduction; Rha S: Isolates from the rhizospheric soil of *Rhanterium epapposum*; Far S: Isolates from the rhizospheric soil of *Farsetia aegyptia*; Hal S: Isolates from the rhizospheric soil of *Haloxylon salicornicum*; Ac, ACN, LTN: Isolates from two set of *Vachellia* *pachyceras* root nodules from KSRI); ARA acetylene reduction assay.

For the primary screening of diazotrophic bacteria, nitrogen free Malate media with bromothymol blue (BTB) was used as an indicator for potential N_2_-fixers. In this process, the diameter for the blue colored producing zone of each bacterial isolate were recorded. About 70, 77, and 73 % of the bacterial isolates were isolated from the rhizospheric soil of *Rhanterium epapposum, Farsetia aegyptia*, and *Haloxylon salicornicum*, respectively, and were found positive for potential nitrogen fixer strains through primary screening. However, around 38% of the bacterial isolates from the root nodule of *Vachellia pachyceras* were found positive for potential nitrogen-fixer strains. Various diameter of colorization zones were recoded from the bacterial isolates ranged from 0.1 cm to 3.6 cm (Table 1).

### Acetylene Reduction Assay (ARA)

When the strains were grown in modified Fraquil agar medium, about 78% of the bacterial isolates were able to grow efficiently on the modified fraquil medium among the primary screened isolates (Figs 1-4, Table 1). The diazotrophic nature of the primary screened isolates were determined by ARA. About 39 isolates exhibited N_2_-fixation ability, but the level of ethylene production (C_2_H_4_) ability of the isolates detected by ARA as positive for N_2_-fixers varied considerably with different isolates (Figs 1-4) and was characterized as low maximum, moderate maximum, high maximum, and very high maximum N_2_-fixation ability (Table 1). However, no direct relationship was established between the diameter of blue coloring zone measured in primary screening and the various levels of N_2_-fixation potential detected by ARA. The results suggest that ARA is the most reliable method of evaluating nitrogen fixation ability of N_2_-fixers. The ARA not only established nitrogenese activity of the bacteria, but also permitted quantification of the nitrogen fixation activity. The amount of ethylene produced in ARA system is shown in the terms of nano moles of ethylene (C_2_H_4_) produced per hour related to cell density measured per milliliter. In the ARA tests, the level of bacterial N_2_-fixation related to conversion of acetylene (C_2_H_2_) to ethylene (C_2_H_4_) varied with different isolates and incubation time, ranged among the strains associated with different rhizospheric soils and root samples from 7 to 145 nmol/h/OD (Figs 1-4). As a matter of comparison, the ethylene production rates of the well-known free-living N_2_ fixer *Azotobacter vinelandii* grown and assayed under similar conditions is around 500 to 3000 nmol/h/OD.

**Fig 1.**
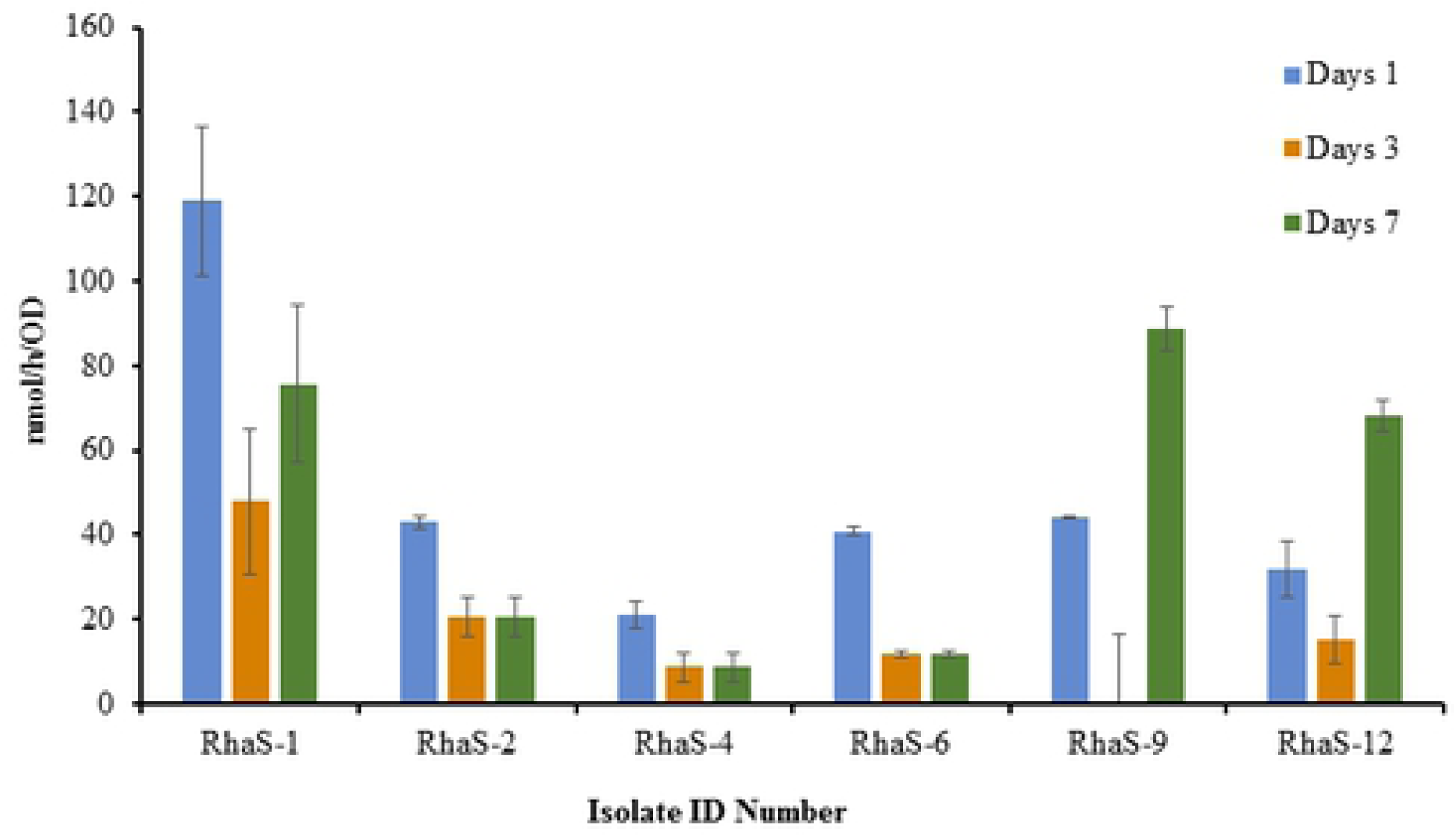
Ethylene production rate (nmol/h) normalized to cell density (OD) measured in the isolate isolated from the rhizospheric soil of *Rhanterium epapposum* after 1, 3, and 7 days of incubation at 28°C in solid medium (modified Fraquil).

**Fig 2.**
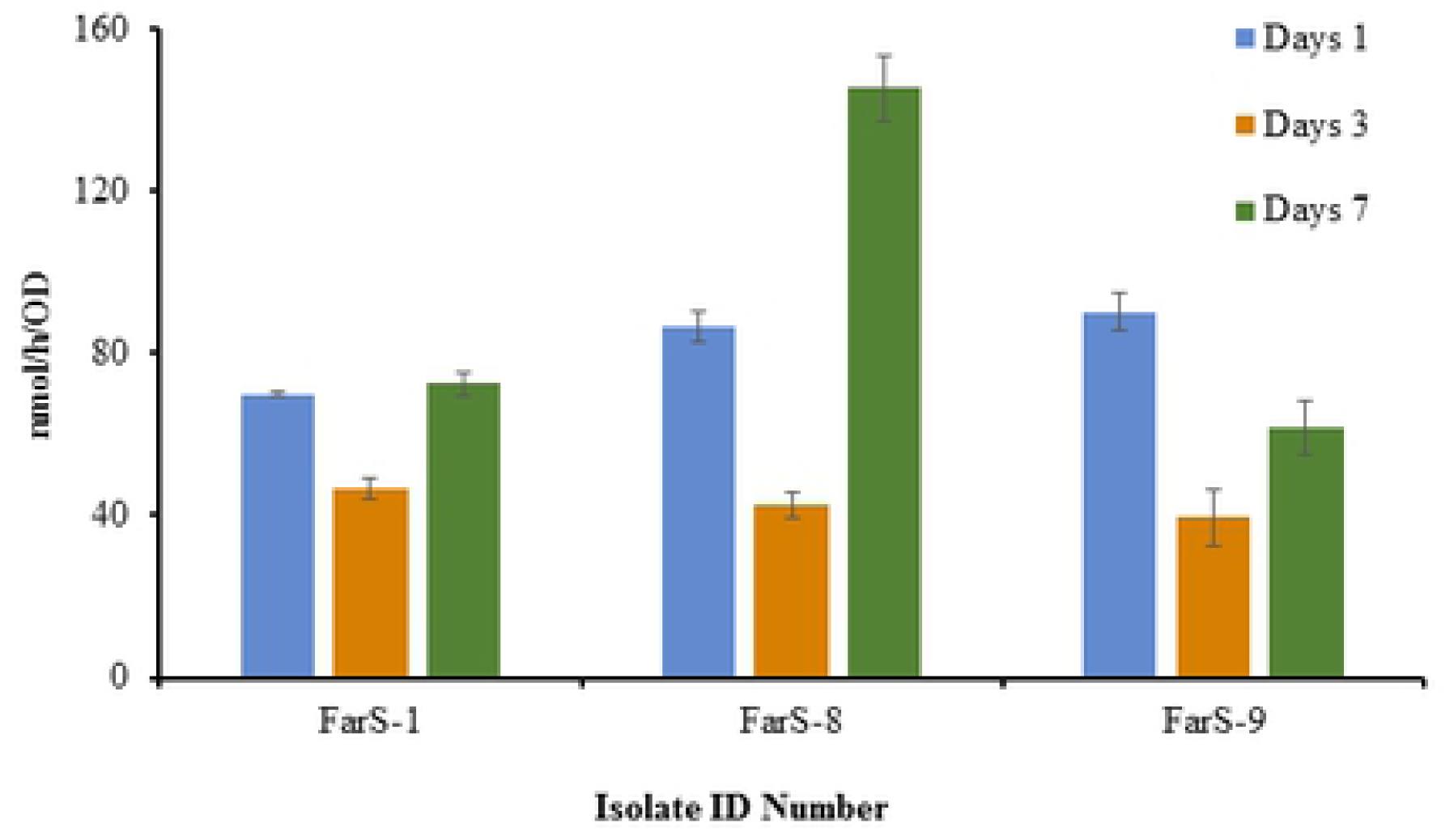
Ethylene production rate (nmol/h) normalized to cell density (OD) measured in the isolate isolated from the rhizospheric soil of *Farsetia aegyptia* after 1, 3, and 7 days of incubation at 28°C in solid medium (modified Fraquil).

**Fig 3.**
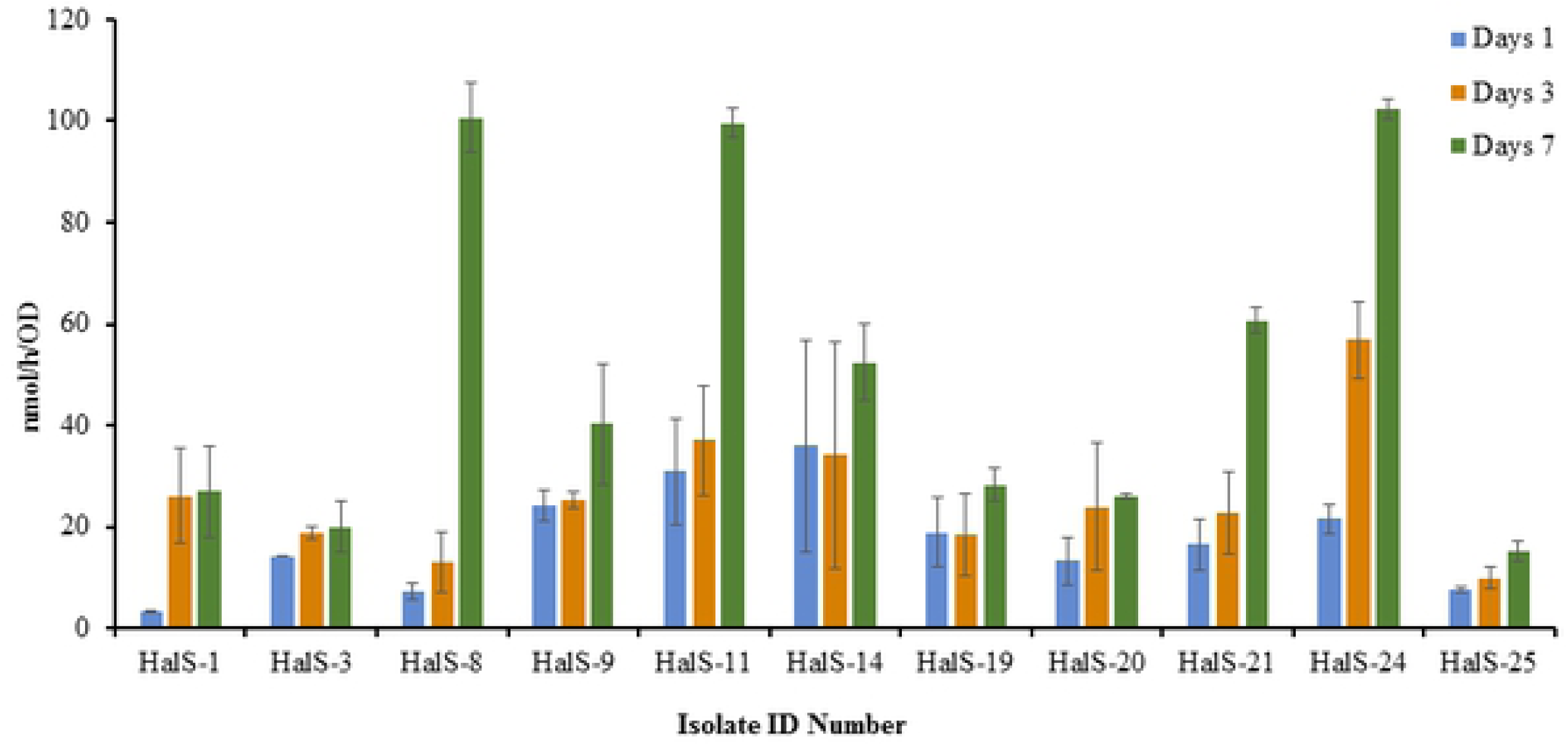
Ethylene production rate (nmol/h) normalized to cell density (OD) measured in the isolate isolated from the rhizospheric soil of *Haloxylon salicornicum* after 1, 3, and 7 days of incubation at 28°C in solid medium (modified Fraquil).

**Fig 4.**
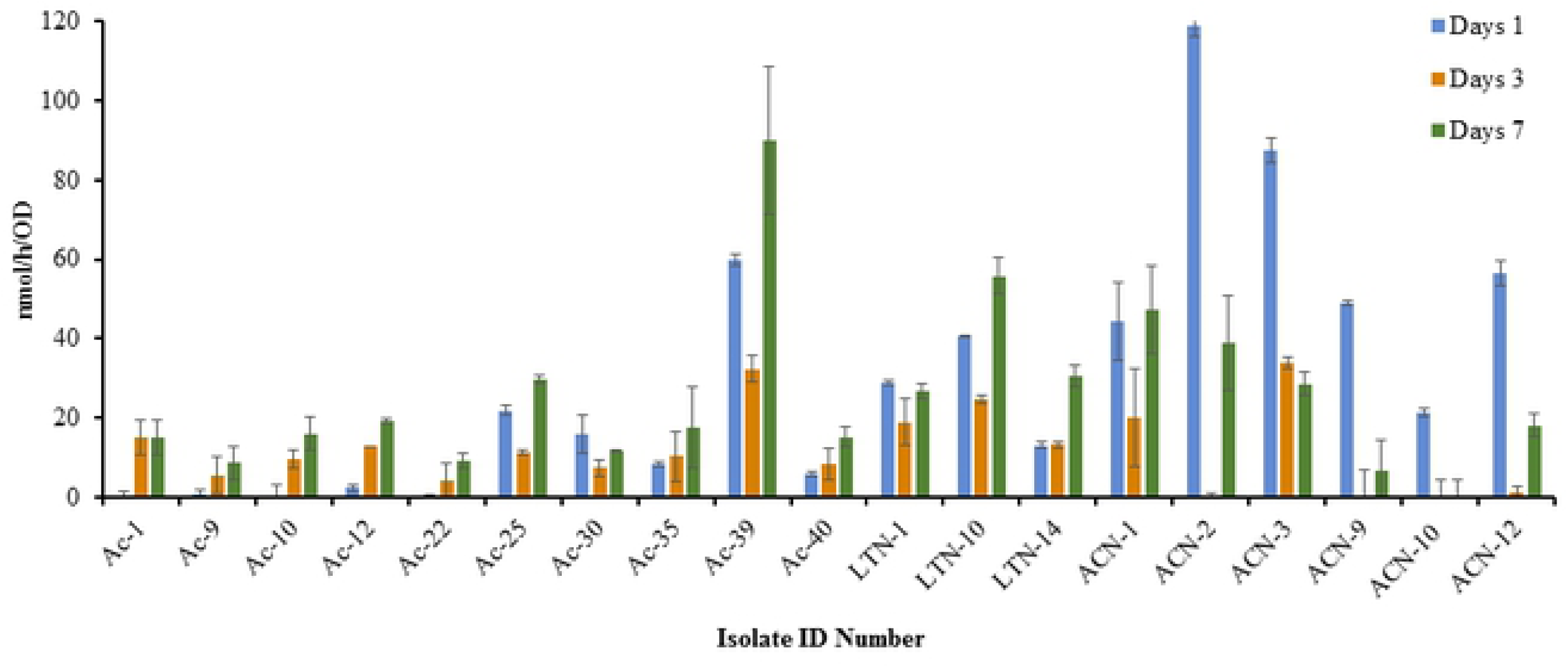
Ethylene production rate (nmol/h) normalized to cell density (OD) measured in the isolate isolated from the root nodule of *Vachellia pachyceras* after 1, 3, and 7 days of incubation at 28°C in solid medium (modified Fraquil).

Interestingly, none of the cultures grew in the liquid modified Fraquil medium. This may be because the cultures were oxygen stressed, due to the agitation (180 rpm) process. Oxygen is a known inhibitor of the enzyme nitrogenase responsible for N_2_ fixation. Static cultures could yield better growth performances. However, these static cultures would likely develop biofilm making bacterial growth difficult to accurately measure. The attempt of screening for N_2-_fixation ability of the isolated strains grown on Yeast Manitol agar and broth media, shows none of the bacterial strain achieved significant N_2_ fixation activity (ARA obtained below detection limit, data not shown). This is likely due to the presence of slight N contamination in the medium.

### Biochemical and molecular identification of the isolates

Of the 50 isolates screened after primary screening, 27 isolates were successfully identified bacterial strains from the rhizospheric soil of *Rhanterium epapposum, Farsetia aegyptia, Haloxylon salicornicum* and root nodules of *Vachellia pachyceras* using BIOLOG® Gen III Microbial Identification System. The isolates belonged to *Rhizobium, Pseudomonas, Bacillus, Enterobacter, Burkholderia, Macroccoccus, Microbactetium, Advenella. Rhizobium* and *Pseudomonas* were the dominant genera with the species *R. radiobacter, R. rhizogenes* among the genera *Rhizobium* and *Pseudomonas stutzeri, Pseudomonas viridilivida, Pseudomonas fluorescens* among the genus *Pseudomonas*. The second dominant genera was *Bacillus* and *Enterobacter* and the species identified were *Bacillus megaterium, Bacillus odyssey, Bacillus simplex/butanolivorans* and *Enterobacter cowanii, Enterobacter homaechei*, respectively. The least dominant species were *Burkholderia anthian/caribensis, Macroccoccus equipercicus, Microbactetium* spp. (CDC.A-4), *Advenella incenata* (Fig 5A). With the biochemical identification system, a single bacterial species *Enterobacter cowanii* was found in the rhizospheric soil of *Rhanterium epapposum*, whereas *Pseudomonas* spp., *Advenella incenata* were found in the rhizospheric soil of *Farsetia aegyptia* and *Pseudomonas stutzeri, Macroccoccus equipercicus* were found in the rhizospheric soil of *Haloxylon salicornicum*. The different bacterial species observed in the root nodules of *Vachellia pachyceras are Rhizobium radiobacter, Rhizobium rhizogenes, Pseudomonas viridilivida, Pseudomonas fluorescens, Bacillus megaterium, Bacillus odyssey, Bacillus simplex*/*butanolivorans, Burkholderia anthian*/*caribensis, Enterobacter homaechei, Microbactetium* spp. (CDC.A-4). Out of 50 isolates tested for biochemical analysis for possible identification, about 23 isolates were unable to be identified and therefore marked as “NO ID” (Table 2, Fig 5A). The reason for not identifying all these isolates could be the new strains or not all the isolates may be supported by the current BIOLOG® database.

**Table 2.**
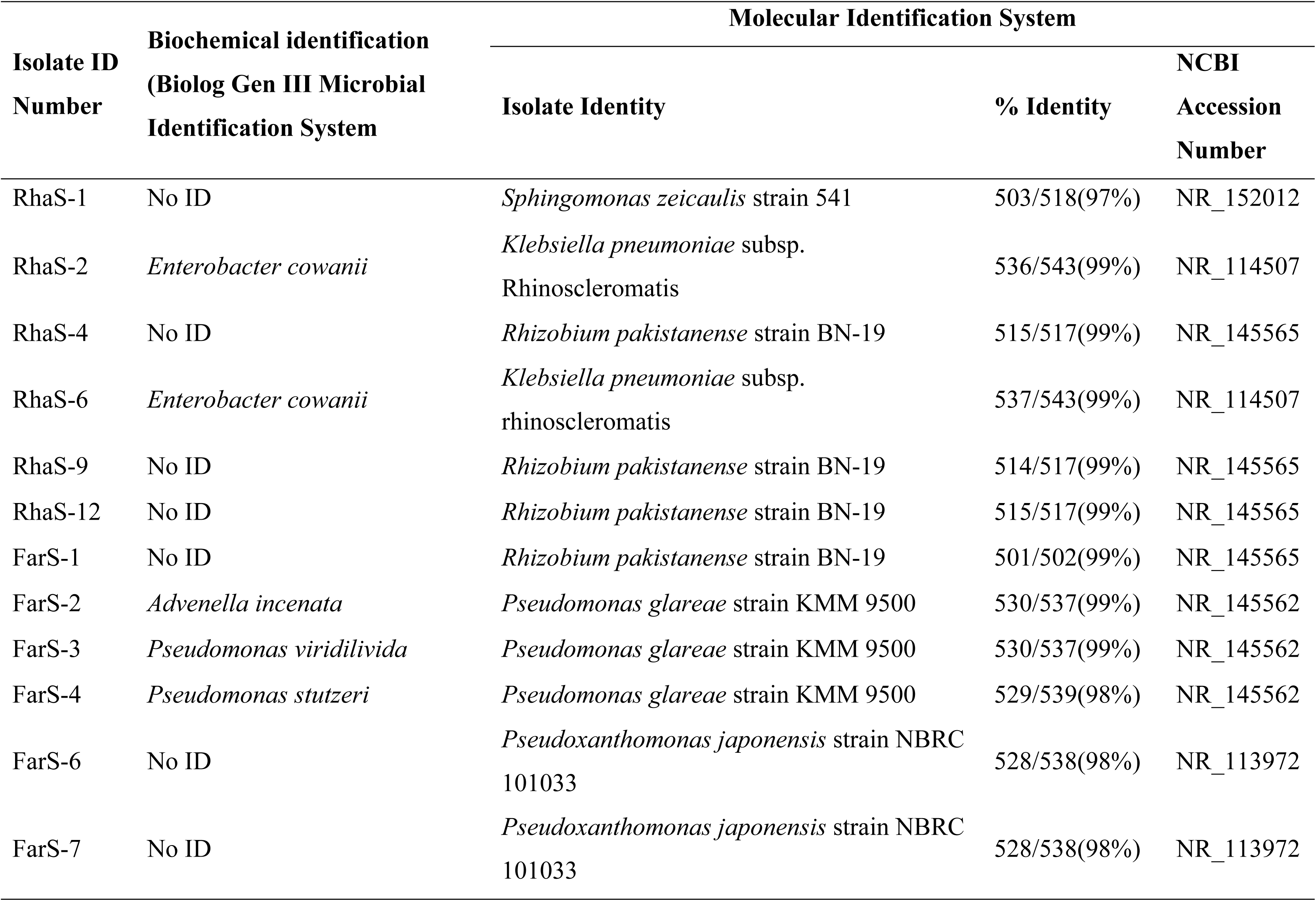
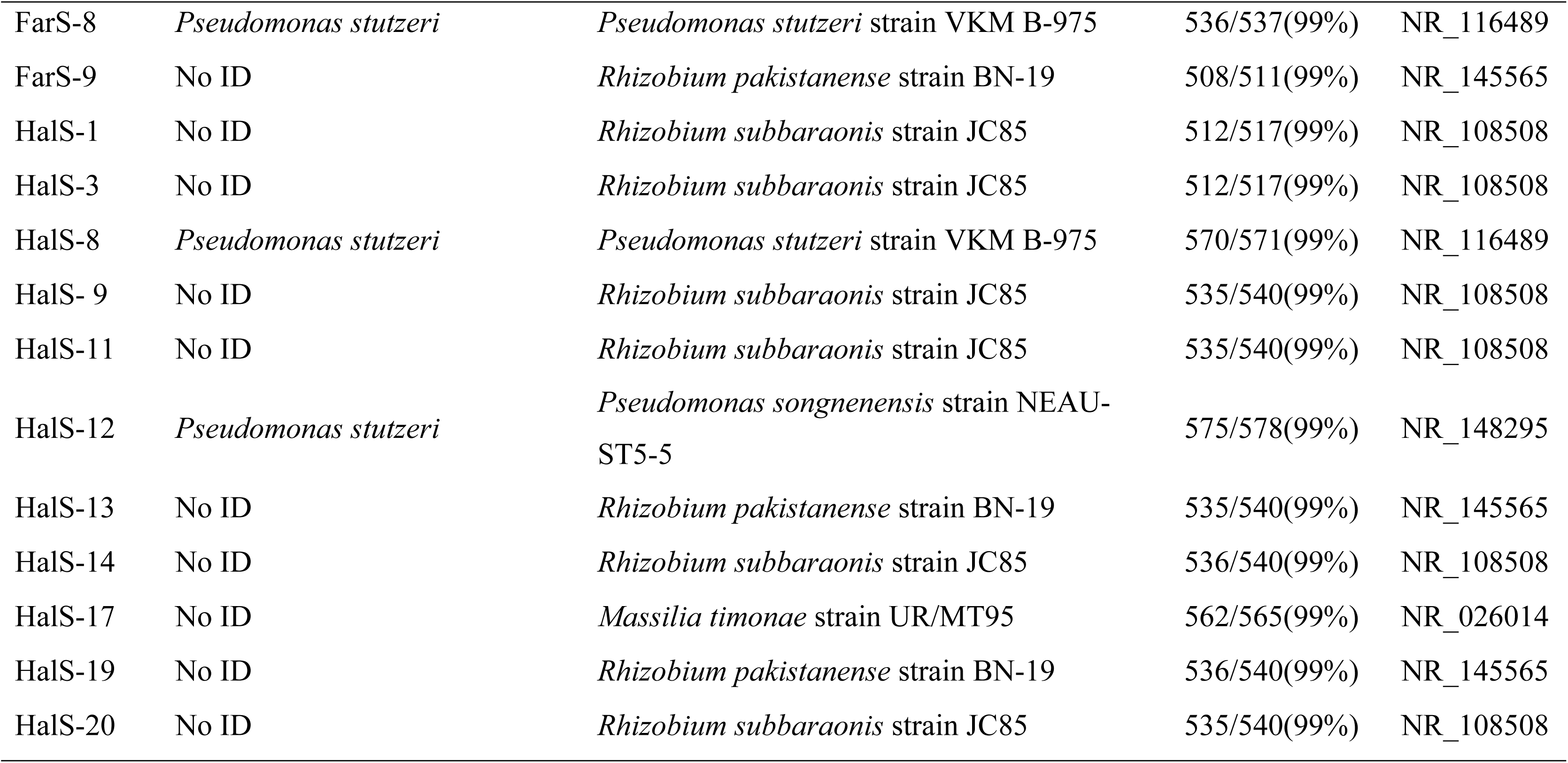
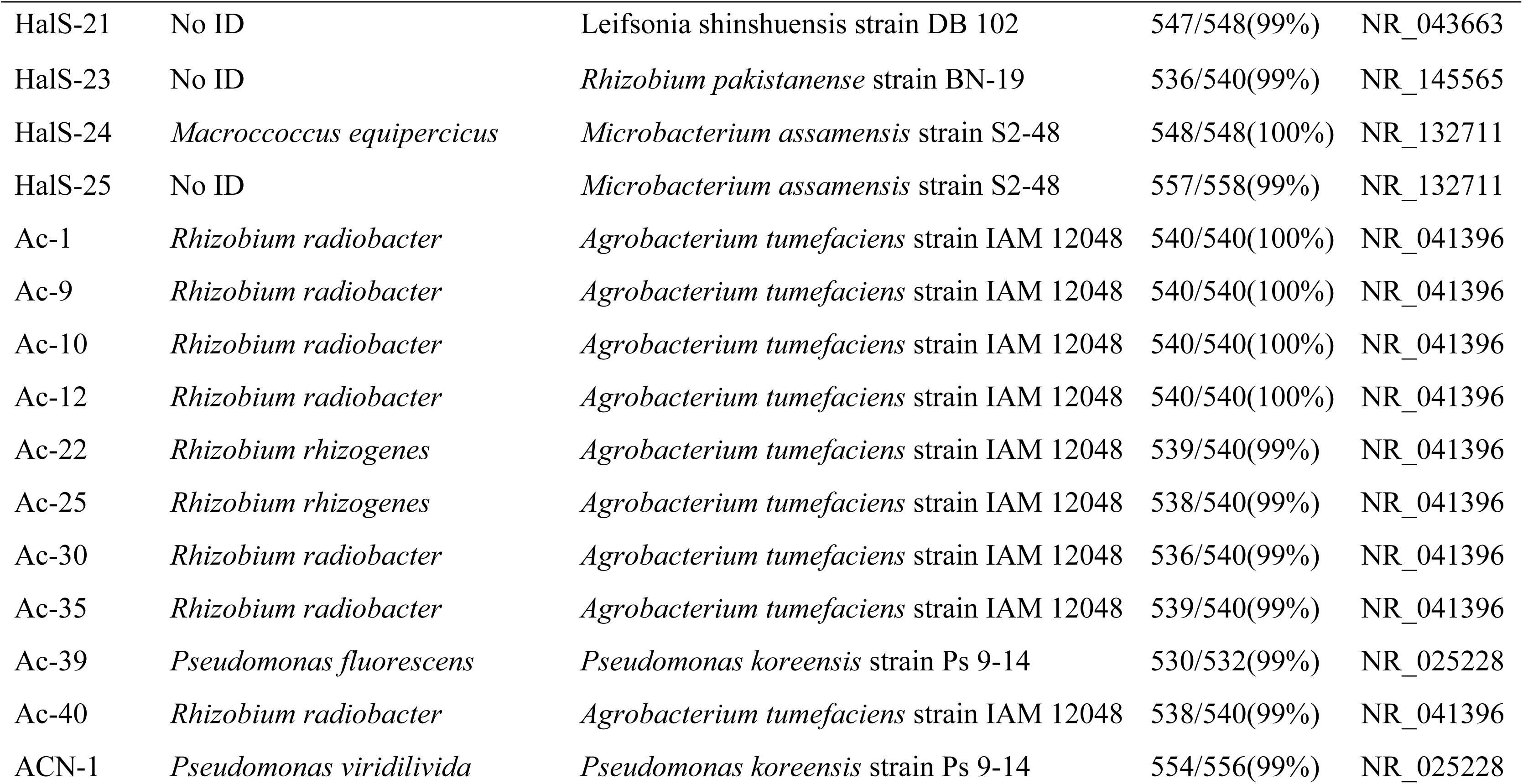
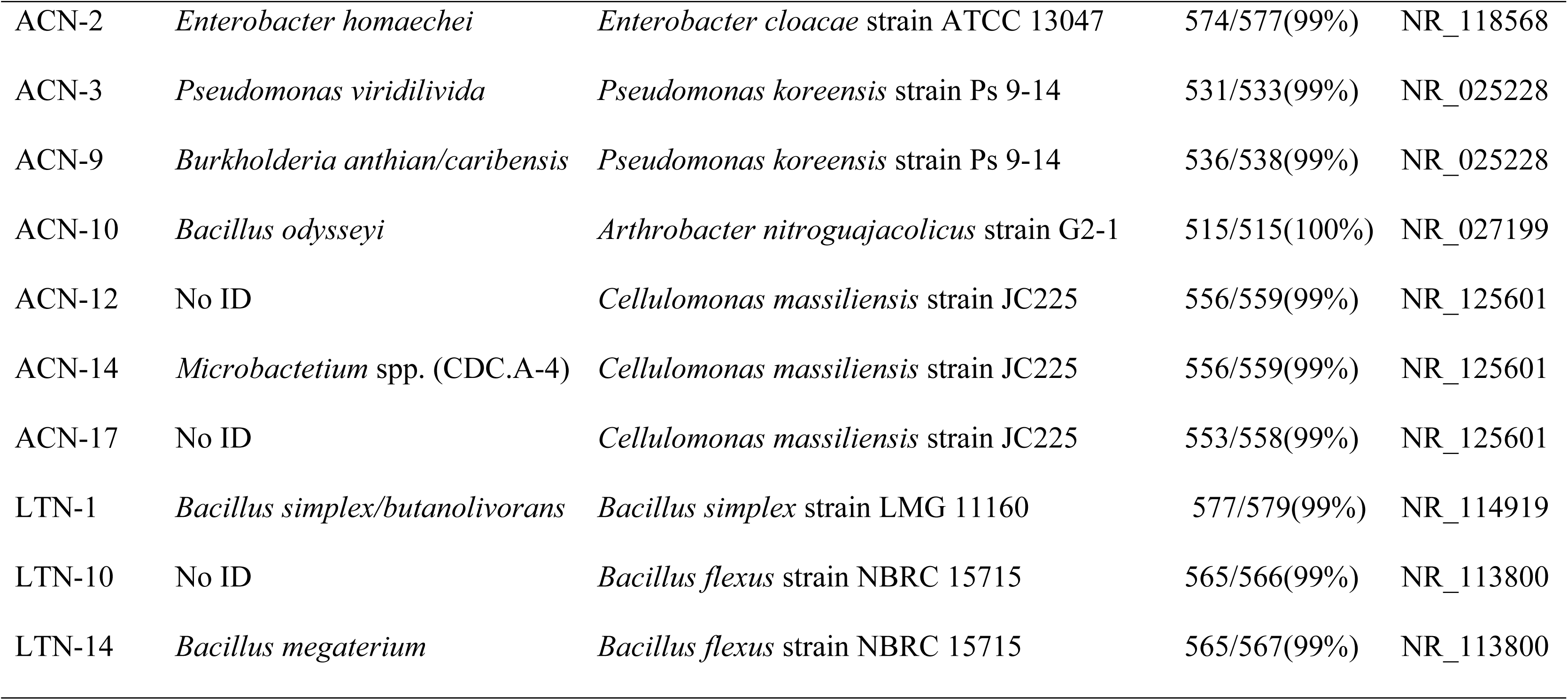
Biochemical and Molecular Identification of the Isolates Positive for Primary Screening for Nitrogen Fixation.

**Fig 5.**
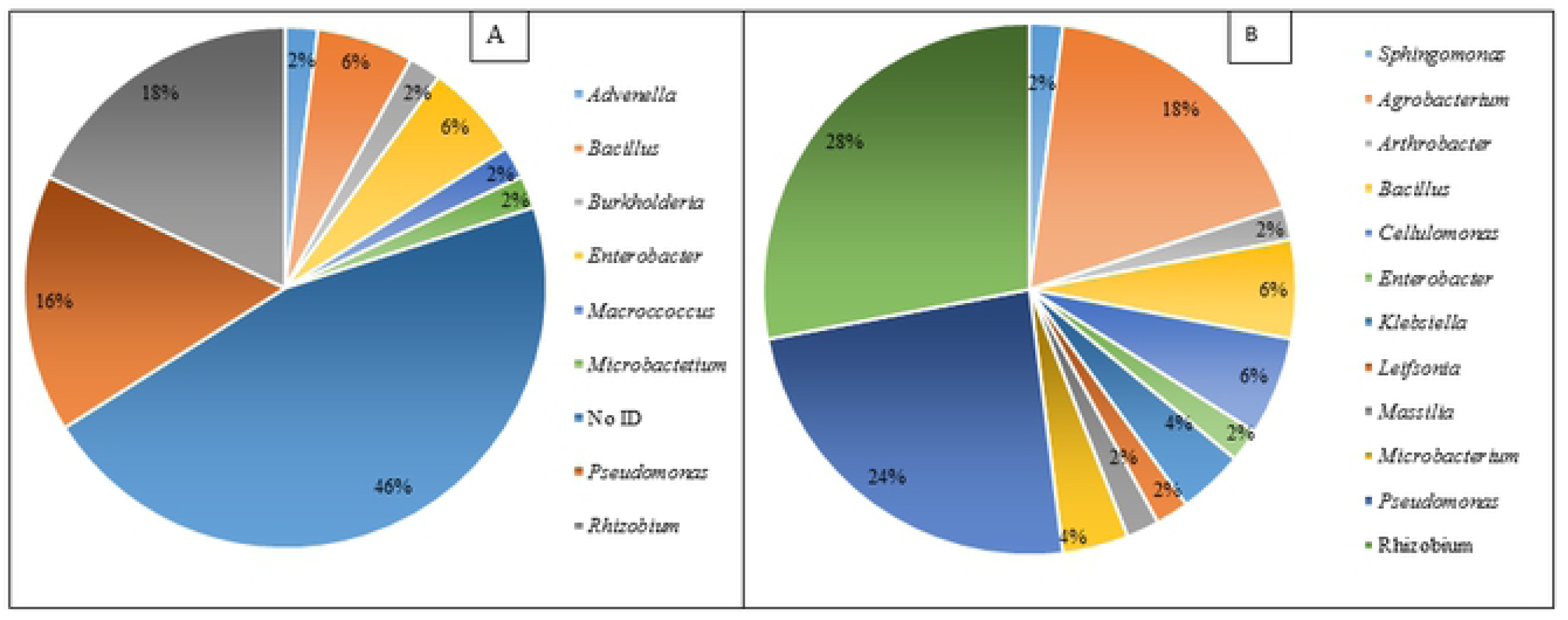
Dominant bacterial genera identified in this study (A), Biochemical identification using BIOLOG® Gen III Microbial Identification System (B), 16s rRNA sequencing.

Hence, it was decided to confirm the identities of isolated strains using molecular identification system via. 16S rRNA gene sequencing method. Partial 16S rRNA gene sequences were obtained for all 50 bacterial strains isolated from the rhizospheric soil of *Rhanterium epapposum, Farsetia aegyptia, Haloxylon salicornicum* and root nodules of *Vachellia pachyceras.* All the isolates including the strains identified under BIOLOG® Gen III Microbial Identification System were identified up to the species level using 16S rRNA sequencing method (Table 2). The BLASTn results of 16S rRNA sequences identified a diverse group of bacterium and majority of the sequences had more than 99% similarity with closely matching sequences existing in the database of Bacterial 16S Ribosomal RNA RefSeq Targeted Loci Project. Only four isolates exhibited less than 99% similarity, *Pseudoxanthomonas japonensis* strain NBRC 101033 (98%), *Pseudomonas glareae* strain KMM 9500 (98%) similarity; *Sphingomonas zeicaulis* strain 541 (97%). Six genera were identified among the 50 identified bacterial strains with the dominant species *Rhizobium* (28%) followed by *Pseudomonas* (24%), *Agrobacterium* (18%), *Bacillus* (6%), *Cellulomonas* (6%), *Klebsiella* (4%), *Microbacterium* (4%) *Sphingomonas* (2%), *Arthrobacter* (2%), *Enterobacter* (2%), *Leifsonia* (2%), *Massilia* (2%) (Fig 5B).

Phylogenetic analysis was conducted for representative bacterial species from each identified bacterial species. The ML and Bayesian analysis showed five major groups of bacterium belonging to the phylum: Gamaproteobacteria, Alphaprotoebacteria, Actinobacteria, Firmicutes and Betaproteobacteria (Fig 6). Gamaproteobacteria was the most dominant group and clustered with seven bacterial species: *Pseudomonas glareae* strain KMM 9500, *Pseudomonas koreensis* strain Ps 9-14, *Pseudomonas songnenensis* strain NEAU-ST5-5, *Pseudomonas stutzeri* strain VKM B-975, *Klebsiella pneumoniae* subsp. *Rhinoscleromatis, Enterobacter cloacae* strain ATCC 13047, *Pseudoxanthomonas japonensis* strain NBRC 101033. Alphaprpteobacteria and Actinobacteria were the second dominant group and clustered with five and four bacterial species, respectively in each group. In Alphaprpteobacteria, species belonged to *Agrobacterium tumefaciens* strain IAM 12048, *Rhizobium pakistanense* strain BN-19, *Rhizobium subbaraonis* strain JC85, and *Sphingomonas zeicaulis* strain 541; Actinobacteria, species belonged to *Cellulomonas massiliensis* strain JC225, *Microbacterium assamensis* strain S2-48, *Arthrobacter nitroguajacolicus* strain G2-1, and *Leifsonia shinshuensis* strain DB 102. Two species clustered in the phylum Firmicutes and a single species present in the phylum Betaproteobacteria. The current study revealed a good number of both free-living and root associated bacterial strains that are present in Kuwait’s desert soils. For the phylogenetic analyses to identify and position the unknown sequences there was a different branching pattern in the phylogenetic tree between ML and Bayesian analyses at the phylum level and hence it did not affect the interpretation and the Bayesian topology is presented in the supplementary material (S1 Fig).

**Fig 6.**
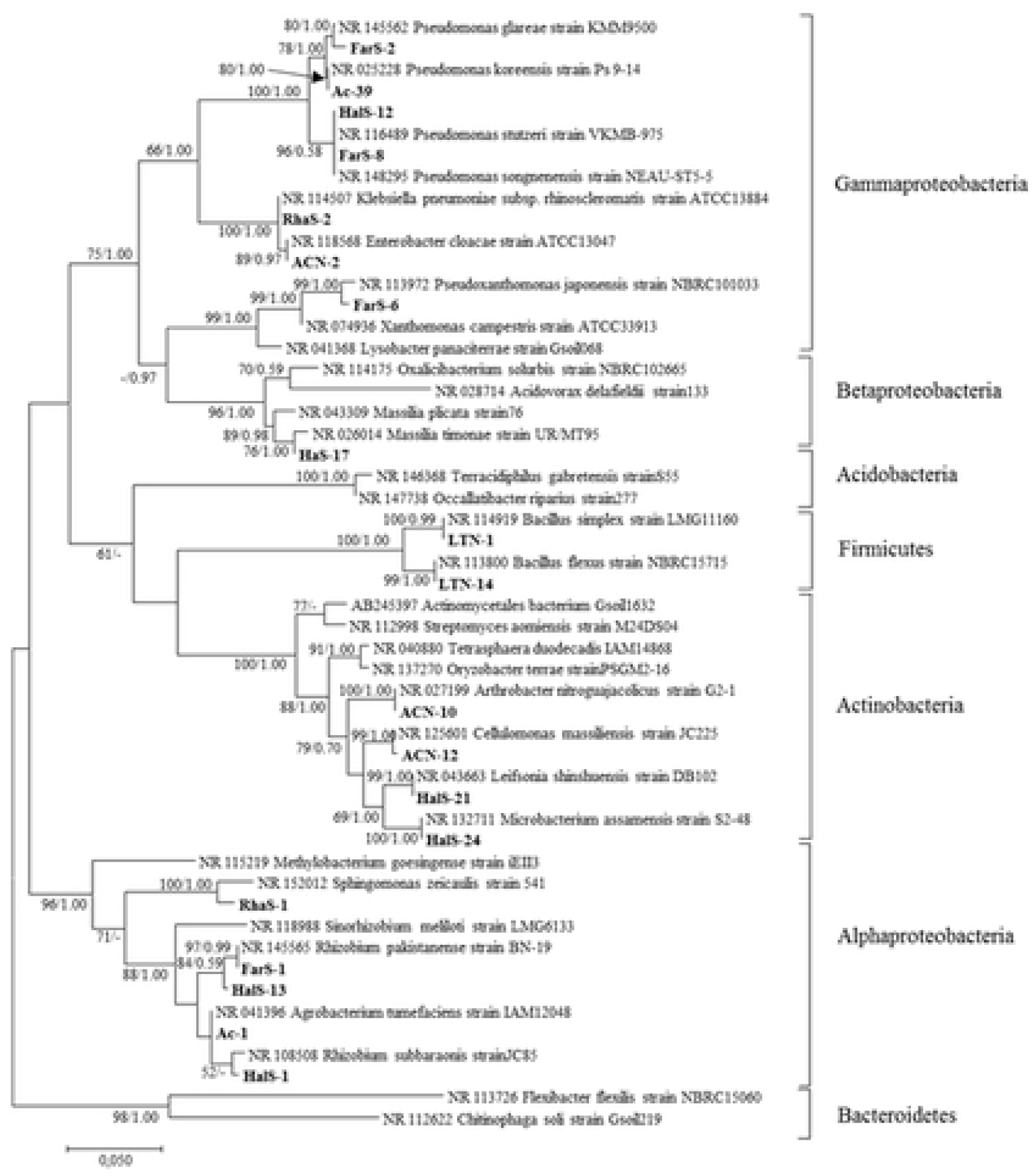
Maximum likelihood analysis of bacteria. Bootstrap percentage values (50 %) generated from 1000 replicates from maximum likelihood and posterior probabilities (>50 %) from Bayesian analysis are shown as [Maximum likelihood bootstrap value/Bayesian posterior probabilities]. Taxa in bold are bacteria from the present study.

## Discussion

The aim of this investigation was to divulge the diazotrophic bacterial community structure associated with the rhizospheric soils of keystone native shrubs and root nodule of a tree species, and to assess their N_2_-fixation ability. According to our knowledge, this study is the first investigation of culture-based isolation, identification, and evaluation of N_2_ -fixing bacteria from rhizospheric soils of Kuwait’s native plant species and report N_2_ fixing ability of isolated bacterial strains using ARA procedure. Available literatures indicate that most of the studies related to N_2_-fixing bacteria were concentrated on symbiosis of different leguminous species. Nevertheless, there is a lack of information about free-living N_2_-fixing bacterial community inhabiting particularly in desert soils, which are able to perform significant N_2_-fixation in desert soils.

Kuwait desert soils are similar to other desert ecosystems characterized by extremely harsh environmental conditions with extreme temperatures, high soil salinity, very low organic matter and nutrients level, high UV radiation levels and instable soil conditions due to dust storms [35]. Despite these prevailing conditions in Kuwait, the current study revealed quite a good diversity of both free-living and root associated bacterial strains in the Kuwait’s desert soils. In this study, free-living diazotrophs were detected in all rhizospheric soils and roots tested. About 50 isolates (potential nitrogen-fixers) detected from primary screening were further tested for biochemical analysis for potential identification. However, only a total of 27 isolates were able to be identified using BIOLOG® (Table 2). The reason for not identifying all the isolates obtained in primary screening could not be ascertain. It is anticipated that there may be the presence of new strains or not all the isolates may be supported by the current BIOLOG® database. However, all the 50 isolates detected from primary screening were identified using 16S rRNA gene sequencing. Phylogenetic analysis based on 16S rRNA sequences of the representative isolates revealed five phyla represented with γ-Proteobacteria, α-Proteobacteria and Actinobacteria being most dominant and followed by, Firmicutes, and β-Proteobacteria (Fig 6). Despite Kuwait desert soils are characterized as hostile environmental conditions, our study identified a great diversity and relatively high abundance of potential nitrogen-fixing bacterial community compared to other ecosystems. There are no previous reports available on isolation and characterization of free-living or entophytic N_2_-fixers from the desert soils of Kuwait. Furthermore, it is interesting to note that the identified bacterial species varied considerably among the rhizospheric soils of different shrubs and root nodules of *Vachellia pachyceras*, suggesting plant species and their rhizospheric effects are important drivers for specificity of microbial diversity in arid soils. Our view on this observation is in agreement with several recent studies that acknowledged plants and root exudates are important driver for the functional gene pool diversity and obtained confirmation of the plant specificity of the diazotrophic communities related to the plants species [1, 36– 38]. Similarly, a significant difference in the free-living nitrogen-fixers was observed due to the type of plant species and suggested as dominant factors determining the structure of diazotrophs [39]. In another recent study by Eida [14], in which the study evaluated the growth promoting properties and salinity tolerance of the bacterial communities associated with the soil, rhizosphere and endosphere of four key desert plant species in Saudi Arabia. The study also revealed a dominance of Actiniobacteria and Protobacteria in all samples tested. Although microbial communities are recruited by plants are dependent by number of factors, such as the genotype of the host plant [40, 41], the crucial determinant of the root-associated bacterial community was recognized as the soil type and the genotype regarded as secondary factor in the root microbial composition [42, 43]. However, in our study we observed varied diazotrophic bacterial composition related to plant species and supports the recent findings [1, 39].

In this current study, among the isolated bacterial strains tested positive from primary screening, about 11 isolates failed to grow in actual nitrogen free media used for ARA. It was difficult to predict why some of these strains did not grow on the modified Fraquil medium. Apparently, some of the cultures may have been affected during the storage period and suggesting repetition of this procedure with use of fresh cultures for confirmation in future studies. The experience in isolating N_2_-fixers from environment suggests that the N_2_-free selective medium used should be maintained pure N_2_-free, any contamination may lead to misleading conclusions and interrupts ARA, which happened in this case, and the isolates were re-grown in absolutely N_2_-free modified fraquil medium for obtaining success in ARA. Our study successfully identified all the 50 nitrogen fixers isolated initially using 16s rRNA gene sequencing, and out of that 78% were confirmed as nitrogen fixers using ARA. Among them, most of the species were from free-living *Rhizobium, Pseudomonas*, and *Agrobacterium* genera followed by *Cellulomonas* and *Bacillus* genera. A great diversity of diazotrophs detected in all the samples tested from desert environment, therefore indicating immense importance of free-living and root associated bacteria in harsh desert conditions and suggesting a great ecological importance of diazotrophs in desert ecosystem

The bacterial conversion of acetylene (C_2_H_2_) to ethylene (C_2_H_4_) production rates of different strains tested in this study are somewhat a lower end compared to the ideal well-known isolates. However, the results are indeed comparable and similar to the studies conducted by Gothwal [22] and Kifle [44] on the diazotrophic bacteria isolated from rhizospheric soils of some important desert plants and maize seedlings. Xu [45] examined putative N_2_-fixing bacteria isolated from rhizospheric soil, root, and stem samples from switchgrass and giant reed and tested for their N2-fixing ability intended to be used as potential biofetilizers. In this study, the levels of bacterial ethylene (C_2_H_4_) production rate in ARA ranged among the strains from 40 to 350 nmol C_2_H_4_ 24 h^-1^ ml^-1^, which are comparable to our study results. Moreover, acetylene reduction in the ARA test and nitrogenase activity level may also be depend on several environmentally and genetically induced factors, such as duration of incubation period and characteristics of strains. Interestingly, most of the isolated rhizobacterial strains (except Ac 39) from root nodules of *Vachellia pachyceras* (Ac) showed a low maximum level of nitrogen fixation potential through acetylene reduction assays compared to all the other strains isolated for this study. The reason for the observed low level of N_2_ fixation potential for the afore-mentioned strains could not be confirmed under the scope of this investigation. However, the bacterial strains associated with the root nodules of *Vachellia pachyceras* (Ac) identified as *Agrobacterium tumefaciens*, which may not be viewed as well known diazotrophs.

A curious observation revealed during identification of isolated bacterial strains (Table 2) using molecular techniques. Unexpectedly, the rhizobacterial strains isolated from root nodules of *Vachellia pachyceras* (Ac) from Sulaiyabia, which were initially identified as *Rhizobium radiobacter* in Biolog Gen III Microbial Identification System and recorded as positive for N_2_-fixation initially by plate test, was interestingly identified as *Agrobacterium tumefaciens* (Table 2) under molecular identification. Several reports indicated that the strain *Agrobacterium tumefaciens* is synonym of *Rhizobium radiobacter* [46]. Indeed this is a noteworthy observation and further investigation is necessary to confirm if *Agrobacterium tumefaciens* may also have the ability to fix atmospheric nitrogen under desert environment. The observation from this study also indicates that biochemical test (Biolog®) for microbial identification may not be the most reliable test for authentic identification.

*Agrobacterium tumefaciens* is a gram-negative bacterium belonging to the family Rhizobiaceae and known to produce crown gall disease in roots of many plants in natural environment. *A. tumefaciens* can live freely in soils, root surface, and inside the plants as parasite. In this investigation, a structure resembles to root nodule often found in Leguminaceae plant roots was observed and assumed similar to typical nitrogen-fixing root nodules produced by *Rhizobium* sp. A few literatures reported that although *A. tumefaciens* is known as pathogenic bacterium for crown gall formation in roots, this bacterium was also found as similar to diazotrophs and report that *A. tumefaciens* can fix nitrogen and are capable of growing on nitrogen-free medium [47, 48]. However, the our finding is the first report as per current knowledge that shows *A. tumefaciens* can produce some sort of root nodules type structure with *Vachellia pachyceras* roots and have the ability to grow on nitrogen-free medium and reduce acetylene to ethylene at low level. The current study recommends further investigation and full understanding of root nodule formation ability and nitrogen-fixating capacity of this bacterium isolated from both rhizospheric soils and root structure. Further research can bring some understanding on the transformation of pathogenic nature of this bacterium to possible nitrogen-fixing and symbiotic relationships with host.

## Conclusion

The current investigation demonstrated successful isolation of free-living N-fixing bacterial strains from rhizospheric soils of three important native shrubs and N-fixers from root nodules of native plant species, *Vachellia pachyceras*. Free-living N-fixing bacteria were found in all rhizospheric soil samples tested. Our study effectively identified all the 50 nitrogen fixers isolated initially using 16s rRNA gene sequencing, and out of that 78% were confirmed as nitrogen-fixers using ARA. Among them, most of the species were from *Rhizobium, Pseudomonas*, and *Agrobacterium* genera followed by *Cellulomonas* and *Bacillus* genera. Further research is needed with these identified isolates to test their effectiveness in nitrogen-fixing ability under desert environment. The current study also warrants additional in-depth research to confirm nitrogen-fixating capacity of *Agrobacterium tumefaciens* isolated from root structure of *Vachellia pachyceras* under desert environment.

## Acknowledgement

The authors are thankful for the constant cooperation and support provided by the management, other departments of KISR and also the assistance provided by the laboratory and field helpers with this research. The authors also acknowledge the technical assistance from the molecular research team at the Université Laval, Québec, Canada for the molecular identification of bacterial strains and Département de chimie, Université de Sherbrooke for acetylene reduction assay.

## Author contributions

**Conceptualization:** M. K Suleiman, A. M. Quoreshi

**Formal analysis:** A. J. Manuvel, M. T. Sivadasan

**Funding acquisition:** M. K Suleiman, A. M. Quoreshi, N. R. Bhat

**Investigation:** M. K Suleiman, A. M. Quoreshi, N. R. Bhat, A. J. Manuvel, M. T. Sivadasan

**Methodology:** M. K Suleiman, A. M. Quoreshi, N. R. Bhat, A. J. Manuvel, M. T. Sivadasan

**Project administration:** M. K Suleiman

**Supervision:** A. M. Quoreshi, M. K Suleiman

**Validation:** M. K Suleiman, A. M. Quoreshi

**Writing –original draft:** A. M. Quoreshi, M. K Suleiman, A. J. Manuvel

**Writing – review & editing:** A. M. Quoreshi, M. K Suleiman, A. J. Manuvel

## Supporting information

**S1 Fig. Bayesian topology of bacterial isolates supporting the branching pattern of identified bacterial isolates (Word)**

## References

1. Köberl J, Prettenthaler F, Bird DN. Modelling climate change impacts on tourism demand: A comparative study from Sardinia (Italy) and Cap Bon (Tunisia). Sci Total environ. 2016; 543: 1039–1053.

2. Hsu SF, Buckley DH. Evidence for the functional significance of diazotroph community structure in soil. The ISME Journal. 2009; 3: 124–136. doi:10.1038/ismej.2008.82.

3. Martínez-Hidalgo P, Hirsch AM. The nodule microbiome: N2-fixing rhizobia do not live alone. Phytobiomes. 2017; 1: 70–82. https://doi.org/10.1094/PBIOMES-12-16-0019-RVW.

4. Gulati A, Sood S, Rahi P, Thakur P, Chauhan S, Chawla I. Diversity Analysis of Diazotrophic Bacteria Associated with the Roots of Tea (Camellia sinensis (L.) O. Kuntze). J Microbiol Biotechn. 2011; 21(6): 545–55. doi: 10.4014/jmb.1012.12022.

5. Simon HM, Smith MW, Herfort L. Metagenomic insights into particles and their associated microbiota in a coastal margin ecosystem. Front Microbiol. 2014; 5: 466. https://doi.org/10.3389/fmicb.2014.00466.

6. Pajares S, Bohannan BJM. Ecology of nitrogen fixing, nitrifying, and denitrifying microorganisms in tropical forest soils. Front Microbiol. 2016; 7: 1045. doi: 10.3389/fmicb.2016.01045.

7. Dixon R, Kahn D. Genetic regulation of biological nitrogen fixation. Nat Rev Microbiol. 2004; 2(8): 621–31. doi: 10.1038/nrmicro954.

8. Ladha JK, Reddy PM. 2000. Steps towards nitrogen fixation in rice: Quest for nitrogen fixation in rice. Proceedings of the 3rd Working Group Meeting on Assessing Opportunities of Nitrogen Fixation in Rice, 2000; August 9-12, 1999, Los Banos, Phillipines, pp: 33-46.

9. Rondon MA, Lehmann J, Ramírez J, Hurtado M. Biological nitrogen fixation by common beans (Phaseolus vulgaris L.) increases with bio-char additions. Biol Fertil Soils. 2007; 43: 699–708. doi 10.1007/s00374-006-0152-z.

10. de Souza R, Beneduzi A, Ambrosini A, Costa PB, Meyer J, Vargas LK. et al. The effect of plant growth-promoting rhizobacteria on the growth of rice (Oryza sativa L.) cropped in southern Brazilian fields. Plant Soil. 2013; 366: 585–603. doi 10.1007/s11104-012-1430-1.

11. Estrada GA, Baldani VID, de Oliveria DM, Urquiaga S, Baldani JI. Selection of phosphate solubilizing diazotrophic Herbaspirillum and Burkholderia strains and their effect on rice crop yield and nutrient uptake. Plant Soil. 2013; 369: 115. https://doi.org/10.1007/s11104012-1550-7.

12. Köberl M, Müller H, Ramadan EM, Berg G. Desert farming benefits from microbial potential in arid soils and promotes diversity and plant health. PLoS ONE. 2011; 6(9): e24452. doi:10.1371/journal.pone.0024452.

13. Abdal MS, Suleiman MK. Soil conservation as a concept to improve Kuwait environment.m Journal Archives of Nature Conservation and Landscape Research. 2002; 41(3-4): 125–130. https://doi.org/10.1080/0003930022000043419.

14. Eida AA, Ziegler M, Lafi FF, Michell CT, Voolstra CR, Hirt H, et al. Desert plant bacteria reveal host influence and beneficial plant growth properties. PLoS ONE. 2018; 13(12): e0208223. https://doi.org/10.1371/journal.pone.0208223.

15. Ortiz N, Armada E, Duque E, Roldán A, Azcón R. Contribution of arbuscular mycorrhizal fungi and/or bacteria to enhancing plant drought tolerance under natural soil conditions: effectiveness of autochthonous or allochthonous strains. J plant physiol. 2015; 174: 87–96. https://doi.org/10.1016/j.jplph.2014.08.019.

16. Friesen ML, Porter SS, Stark SC, von Wettberg EJ, Sachs JL, Martinez-Romero E. Microbially mediated plant functional traits. Annu Rev Ecol Evol S. 2011; 42: 23–46. https://doi.org/10.1146/annurev-ecolsys-102710-145039.

17. de Zelicourt A, Al-Yousif M, Hirt H. Rhizosphere microbes as essential partners for plant stress tolerance. Mol plant. 2013; 6(2): 242–5. doi:10.1093/mp/sst028.

18. Mengual C, Schoebitz M, Azcón R, Roldán A. Microbial inoculants and organic amendment improves plant establishment and soil rehabilitation under semiarid conditions. J environ manage. 2014; 134: 1–7. https://doi.org/10.1016/j.jenvman.2014.01.008.

19. Kieft T, Skujinš J. Soil microbiology in reclamation of arid and semiarid lands. In: Skujins J, editor. Semiarid lands and deserts: soil resource and reclamation. New York: Marcel Dekker; 1991. p. 209–56.

20. Eckford, R, Cook FD, Saul D, Aislabie J, Foght J. Free-living heterotrophic nitrogen-fixing bacteria isolated from fuel-contaminated Antarctic soils. Appl Environ Microbiol. 2002; 68(10): 5181–5185. doi: 10.1128/AEM.68.10.5181-5185.2002.

21. KüçükÇ, M. Kivanç M, Kinaci E. Characterization of Rhizobium sp. isolated from bean. Turk J Biol. 2006; 30: 127–132.

22. Gothwal, RK, Nigam VK, Mohan MK, Sasmal D, Ghosh P. Screening of nitrogen fixers from rhizospheric bacterial isolates associated with important desert plants. Appl Ecolo Env Res. 2007; 6(2): 101–109.

23. Muyzer G, de Waal EC, Uitterlinden AG. Profiling of complex microbial populations by denaturing gradient gel electrophoresis analysis of polymerase chain-amplified gene coding for 16S rRNA. Appl Environ Microbiol. 1993; 59: 695–700.

24. 154. Lane DJ, Pace B, Olsen GJ, Stahl DA, Sogin ML, Pace NR. Rapid determination of 16S ribosomal RNA sequences for phylogenetic analyses. Proc Natl Acad Sci USA. 1985; 82: 6955–6959. https://doi.org/10.1073/pnas.82.20.6955.

25. Hall T. Biological sequence alignment editor for Win95/98//NT/2K/XP. Available from: http://www.mbio.ncsu.edu/BioEdit/bioedit.html

26. Hall TA. BioEdit: a user-friendly biological sequence alignment editor and analysis program for Windows 95/98/NT. Nucleic Acids Symp. 1999; 41: 95–98.

27. BLAST: The Basic Local Alignment Search Tool. Available from: http://www.ncbi.nlm.nih.gov/BLAST/.

28. Thompson K, Bakker JP, Bekker RM. Soil seed banks of north west Europe: methodology, density and longevity. 1997. Cambridge: Cambridge University Press.

29. MEGA: Molecular Evolutionary Genetics Analysis. Available from: http://www.megasoftware.net.

30. Kumar S, Stecher G, Tamura K. MEGA7: Molecular Evolutionary Genetics Analysis Version 7.0 for Bigger Datasets. Mol Biol Evol. 2016; 33(7): 1870–1874. https://doi.org/10.1093/molbev/msw054.

31. Saitou N, Nei M. The neighbor-joining method: A new method for reconstructing phylogenetic trees. Mol Bio Evol. 1987; 4: 406–425. https://doi.org/10.1093/oxfordjournals.molbev.a040454.

32. Kimura M. A simple method for estimating evolutionary rate of base substitutions through comparative studies of nucleotide sequences. J Mol Evol. 1980; 16: 111–120. https://doi.org/10.1007/BF01731581.

33. Felsenstein J. (1985) Confidence limits on phylogenies: An approach using the bootstrap. Evolution. 1985; 39: 783–791. https://doi.org/10.1111/j.15585646.1985.tb00420.x.

34. Ronquist F, Huelsenbeck JP. Mr.Bayes 3: Bayesian phylogenetic inference under mixed models. Bioinformatics. 2003; 19(12): 1572–1574. https://doi.org/10.1093/bioinformatics/btg180.

35. Cary SC, McDonald IR, Barrett JE, Cowan DA. On the rocks: the microbiology of Antarctic dry valley soils. Nat Rev Microbiol. 2010; 8: 129–138. doi:10.1038/nrmicro2281.

36. Bais HP, Weir TL, Perry LG, Gilroy S, Vivanco JM. The role of root exudates in rhizosphere interactions with plants and other organisms. Annu Rev Plant Biol. 2006; 57: 233–66. https://doi.org/10.1146/annurev.arplant.57.032905.105159.

37. Berg G, Smalla K. Plant species and soil type cooperatively shape the structure and function of microbial communities in the rhizosphere. FEMS Microbiol Ecol. 2009; 68: 1–13. https://doi.org/10.1111/j.1574-6941.2009.00654.x.

38. Bulgarelli D, Rott M, Schlaeppi K, van Themaat EVL, Ahmadinejad N, Assenza F, et al. Revealing structure and assembly cues for Arabidopsis root-inhabiting bacterial microbiota. Nature. 2012; 488(7409): 91–5. doi:10.1038/nature11336.

39. Zhan J, Sun Q. Diversity of free-living nitrogen-fixing microorganisms in the rhizosphere and non-rhizosphere of pioneer plants growing on wastelands of copper mine tailings. Microbiol Res. 2012 20;167(3): 157–65. https://doi.org/10.1016/j.micres.2011.05.006.

40. Bulgarelli D, Schlaeppi K, Spaepen S, van Themaat EVL, Schulze-Lefert P. Structure and functions of the bacterial microbiota of plants. Annu Rev Plant Biol. 2013; 64: 807–38. https://doi.org/10.1146/annurev-arplant-050312-120106.

41. Bulgarelli D, Garrido-Oter R, Münch Philipp C, Weiman A, Dröge J, Pan Y, et al. Structure and function of the bacterial root microbiota in wild and domesticated barley. Cell Host & Microbe. 2015; 17(3): 392–403. https://doi.org/10.1016/j.chom.2015.01.011.

42. Lundberg DS, Lebeis SL, Paredes SH, Yourstone S, Gehring J, Malfatti S, et al. Defining the core Arabidopsis thaliana root microbiome. Nature. 2012; 488: 86. doi: 10.1038/nature11237.

43. Yeoh YK, Dennis PG, Paungfoo-Lonhienne C, Weber L, Brackin R, Ragan MA, et al. Evolutionary conservation of a core root microbiome across plant phyla along a tropical soil chronosequence. Nature Communications. 2017; 8. https://doi.org/10.1038/s41467-017-00262-8.

44. Kifle MH, Laing MD. Effects of Selected Diazotrophs on Maize Growth. Front Plant Sci. 2016; 7: 1429. doi: 10.3389/fpls.2016.01429.

45. Xu J, Kloepper JW, Huang P, McInroy JA, Hu CH. Isolation and characterization of N2-fixing bacteria from giant reed and switchgrass for plant growth promotion and nutrient uptake J Basic Microbiol. 2018; 58(5): 459–471. doi: 10.1002/jobm.201700535.

46. Koivunen ME, Morisseau C, Horwath WR, Hammock BD. Isolation of a strain of Agrobacterium tumefaciens (Rhizobium radiobacter) utilizing methylene urea (ureaformaldehyde) as nitrogen source. Can. J. Microbiol. 2004; 50: 167–174. doi: 10.1139/W04-001.

47. Kanvinde L, Sastry GRK. Agrobacterium tumefaciens is a diazotrophic bacterium. Appl Environ Microbiol. 1990; 56(7): 2087–2092.

48. My PT, Manucharova NA, Stepanov AL, Pozdyakov, LA, Selitskaya OV, Emtsev VT. 2015.Agrobacterium tumefaciens as associative nitrogen-fixing bacteria. Moscow University Soil Science Bulletin. 2015; 70(3): 133–138. https://doi.org/10.3103/S0147687415030047.

